# The role of deleterious substitutions in crop genomes

**DOI:** 10.1101/033175

**Authors:** Thomas J. Y. Kono, Fengli Fu, Mohsen Mohammadi, Paul J. Hoffman, Chaochih Liu, Robert M. Stupar, Kevin P. Smith, Peter Tiffin, Justin C. Fay, Peter L. Morrell

## Abstract

Populations continually incur new mutations with fitness effects ranging from lethal to adaptive. While the distribution of fitness effects (DFE) of new mutations is not directly observable, many mutations likely have either no effect on organismal fitness or are deleterious. Historically, it has been hypothesized that a population may carry many mildly deleterious variants as segregating variation, which reduces the mean absolute fitness of the population. Recent advances in sequencing technology and sequence conservation-based metrics for inferring the functional effect of a variant permit examination of the persistence of deleterious variants in populations. The issue of segregating deleterious variation is particularly important for crop improvement, because the demographic history of domestication and breeding allows deleterious variants to persist and reach moderate frequency, potentially reducing crop productivity. In this study, we use exome resequencing of fifteen barley accessions and genome resequencing of eight soybean accessions to investigate the prevalence of deleterious SNPs in the protein-coding regions of the genomes of two crops. We conclude that individual cultivars carry hundreds of deleterious SNPs on average, and that nonsense variants make up a minority of deleterious SNPs. Our approach annotates known phenotype-altering variants as deleterious more frequently than the genome-wide average, suggesting that putatively deleterious variants are likely to affect phenotypic variation. We also report the implementation of a SNP annotation tool (BAD_Mutations) that makes use of a likelihood ratio test based on alignment of all currently publicly available Angiosperm genomes.

## Introduction

Mutation produces a constant influx of genetic variants into populations. Each mutation has a fitness effect that varies from lethal to neutral to advantageous. While the distribution of fitness effects of new mutations is not directly observable (Eyre-Walker and Keightley 2007), most mutations with fitness impacts are deleterious (Keightley and Lynch 2003). It is generally assumed that deleterious mutations alter phylogenetically-conserved sites (Doniger et al. 2008), or cause loss of protein function (Yampolsky et al. 2005). Strongly deleterious mutations (particularly those with dominant effects) are quickly purged from populations by purifying selection. Likewise, strongly advantageous mutations increase in frequency, and ultimately fix due to positive selection (Robertson 1960; Smith and Haigh 1974). Weakly deleterious mutations have the potential to persist in populations and cumulatively contribute significantly to reductions in fitness as segregating deleterious variants (Fay et al. 2001; Eyre-Walker et al. 2006; Doniger et al. 2008).

Considering a single variant in a population, three parameters affect its segregation: the effective population size (*N_e_*), the selective coefficient against homozygous individuals (*s*), and the dominance coefficient (*h*). The effects of *N_e_* and s are relatively simple; variants are primarily subject to genetic drift rather than selection if *N_e_s* < 1 (Kimura et al. 1963). The effect of *h* is not as straightforward, as it depends on the genotypic frequencies and the degree of outcrossing in the population. In populations with a high degree of self fertilization or sibling mating, many individuals will be homozygous, which reduces the importance of *h* in determining the efficacy of selection against the variant (Glémin 2003). In populations that are closer to panmixia, an individual deleterious variant will occur primarily in the heterozygous state, and *h* will determine how “visible” the variant is to selection, with higher values of *h* increasing the efficacy of selection (Charlesworth and Charlesworth 1999). A completely recessive deleterious variant may remain effectively neutral as long as the frequency of the variant is low enough such that there are not a substantial number of homozygous carriers. Conversely, a completely dominant deleterious variant is expected to be quickly purged from the population (Lande and Schemske 1985). On average, deleterious variants segregating in a population are predicted to be partially recessive (Simmons and Crow 1977), allowing them to remain “hidden” from the action of purifying selection, and reach moderate frequencies. This may be expected, for example, based on data from a gene knockout library in yeast (Shoemaker et al. 1996), which indicate that protein loss-of-function variants have an average dominance coefficient of 0.2 (Agrawal and Whitlock 2012).

Effective recombination rate also has important impacts on the number and distribution of deleterious mutations in the genome. Regions with low effective recombination are prone to the irreversible accumulation of deleterious variants. This phenomenon is known as the “ratchet effect” (Muller 1964). In finite populations with low recombination, the continual input of deleterious mutations and stochastic variation in reproduction causes the loss of individuals with the fewest deleterious variants. Lack of recombination precludes the selective elimination of chromosomal segments carrying deleterious variants, and thus they can irreversibly increase, similar to how a ratchet turns in only one direction (Muller 1964). Nordborg (2000) demonstrates that under high levels of inbreeding, effective recombination rate can be decreased by almost 20-fold relative to an outbreeding population, showing that mating system can be a major determinant in the segregation of deleterious variation. While inbreeding populations are especially susceptible to ratchet effects on a genome-wide scale, even outbreeding species have genomic regions with limited effective recombination (Arnheim et al. 2003; McMullen et al. 2009). In maize, these low recombination regions are observed to harbor excess heterozygosity in inbred lines, suggesting that they maintain deleterious variants that cannot be made homozygous (Ridgers-Melnick et al. 2014). Both simulation studies (Felsenstein 1974) and empirical investigations in *Drosophila melanogaster* (Campos et al. 2012, 2014) indicate that deleterious variants accumulate in regions of limited recombination.

Efforts to identify deleterious variants and quantify them in individuals have led to a new branch of genomics research. In humans, examination of the contribution of rare deleterious variants to heritable disease has contributed to the emergence of personalized genomics as a field of study (reviewed in Abecasis et al. 2010; Cooper et al. 2010; Marth et al. 2011). Current estimates suggest that an average human may carry ~300 loss-of-function variants (Abecasis et al. 2010; Agrawal and Whitlock 2012) and up to tens of thousands of weakly deleterious variants in coding and functional noncoding regions of the genome (Arbiza et al. 2013). In terms of effects on organismal fitness, the average human carries three lethal equivalents (Gao et al. 2015; Henn et al. 2015). These variants are enriched for mutations that are causative for diseases (Kryukov et al. 2007; Marth et al. 2011). As such they are expected to have appreciable negative selection coefficients (*N_e_s*) and be kept at low frequencies due to the action of purifying selection.

Humans are not unique in harboring substantial numbers of deleterious variants. It is estimated that almost 40% of nonsynonymous variants in *Saccahromyces cerevisiae* have deleterious effects (Doniger et al. 2008) and 20% of nonsynonymous variants in rice (Lu et al. 2006), *Arabidopsis thaliana* (Günther and Schmid 2010), and maize (Mezmouk and Ross-Ibarra 2014) are deleterious. In dogs, Cruz et al. (2008) identified an excess of nonsynonymous single nucleotide polymorphisms (SNPs) segregating in domesticated dogs relative to grey wolves. A similar pattern has been found in horses (Schubert et al. 2014) and sunflowers (Renault and Rieseberg 2015), suggesting that an increased prevalence of deleterious variants may be a “cost of domestication.”

Approaches to identify deleterious mutations take one of two forms. Quantitative genetic approaches have been employed that investigate the aggregate impact of potentially deleterious alleles on fitness. Mutation accumulation studies (e.g., Mukai 1964; Schultz et al. 1999; Shaw et al. 2002; Charlesworth et al. 2004) use change in fitness over generations within lineages to estimate mutational effects on fitness. Coupled with DNA sequencing technologies, these studies may shed light on how many DNA sequence changes are potentially deleterious (e.g., Ossowski et al. 2010). On the other hand, purely bioinformatic approaches make use of measures of sequence conservation to identify variants with a significant probability of being deleterious. When combined with genome-scale resequencing, they permit the identification of large numbers of putatively deleterious variants. Commonly applied approaches include SIFT (Sorting Intolerant From Tolerated) (Ng 2003), PolyPhen2 (Polymorphism Phenotyping) (Adzhubei et al. 2010), and a likelihood ratio test (LRT) (Chun and Fay 2009). These sequence conservation approaches operate in the absence of phenotypic data, but allow assessment of individual sequence variants. As such, some variants identified bioinformatically may be locally adaptive, or conditionally neutral. However, given the observation that deleterious mutations constantly arise and continue to segregate in populations, their targeted identification and elimination from breeding populations presents a novel path for crop improvement (Morrell et al. 2011).

In this study, we investigate the distribution of deleterious variants in thirteen barley (*Hordeum vulgare* ssp. *vulgare*) cultivars, two wild barley (*H. vulgare* ssp. *spontaneum*) accessions, seven soybean (*Glycine max*) cultivars, and one wild soybean (*Glycine soja*) accession using exome and whole genome resequencing. We seek to answer four questions about the presence of deleterious variants: *i*) How many deleterious variants do individual cultivars harbor, and what proportion of these are nonsense (early stop codons) versus nonsynonymous (missense) variants? *ii*) What proportion of nonsynonymous variation is inferred to be deleterious? *iii*) How many known phenotype-altering SNPs are inferred to be deleterious? *iv*) How does the relative frequency of deleterious variants vary with recombination rate? We identify an average of ~1,000 deleterious variants per accession in our barley sample and ~700 deleterious variants per accession in our soybean sample. Approximately 40% of the deleterious variants are private to one individual in both species, suggesting the potential for selection for individuals with a reduced number of deleterious variants. Approximately 3-6% of nonsynonymous variants are inferred to be deleterious by all three annotation approaches used in our study, and known causative SNPs annotate as deleterious at a much higher proportion than the genomic average. In soybean, where genome-wide recombination rate estimates are available, the proportion of deleterious variants is negatively correlated with recombination rate.

## Results

### Variant Calling and Identification of Deleterious SNPs

Resequencing and read mapping followed by read de-duplication resulted in an average coverage of ~39X exome coverage for our barley samples and ~38X genome coverage in soybean. A summary of our resequencing data and read mapping statistics is shown in Table S1. Average heterozygosity was 2.5% in our barley sample, and 0% in our soybean sample, reflecting the inbreeding of the accessions. The observed heterozygosity in our barley sample is mostly due to the inclusion of wild material, which is less inbred than the cultivars. Heterozygous variant calls in soybean were all in reads with low mapping score, possibly due to the highly duplicated nature of the soybean genome (Schmutz et al. 2010). A table of the barley accessions used in this study is shown in Table S2, and the soybean accessions are shown in Table S3.

After realignment and variant recalibration, we identified 652,797 SNPs in thirteen cultivated and two wild barley lines. The majority of these SNPs were noncoding, with 522,863 occurring outside of coding sequence (CDS) annotations. Of the coding SNPs, 70,069 were synonymous, and 59,865 were nonsynonymous. A summary of the variants in various functional classes is shown in Table 1, and a per-sample summary is shown in Table S4. SIFT identified 13,626 SNPs as deleterious, PolyPhen2 identified 13,534 SNPs to be deleterious, and the LRT called 17,865 deleterious. The intersection of all three approaches identifies a much smaller set of deleterious variants, with a total of 4,872 nonsynonymous SNPs identified as deleterious. While individual methods identified ~18% of nonsynonymous variants as deleterious, the intersect of approaches identifies 5.7%. A derived site frequency spectrum (SFS) of synonymous, nonsynonymous, and putatively deleterious SNPs in our barley sample is shown in Figure 1A.

**Table 1.** Mean numbers of SNPs in various classes.

| Species | Diff. from Ref. | Noncoding | Syn. | Nonsyn. | Nonsense |
| --- | --- | --- | --- | --- | --- |
| Barley | 162,954<br>(51,231.34) | 115,456<br>(41,065.22) | 15,591<br>(5,691.81) | 12,351<br>(4,492.53) | 77<br>(33.13) |
| Soybean | 82,840<br>(56,780.03) | 44,704<br>(29,477.65) | 14,167<br>(8,161.21) | 18,695<br>(11,289.72) | 540<br>(345.05) |

Syn. = Synonymous; Nonsyn. = Nonsynonymous. Numbers are Mean (sd)

**Figure 1:**
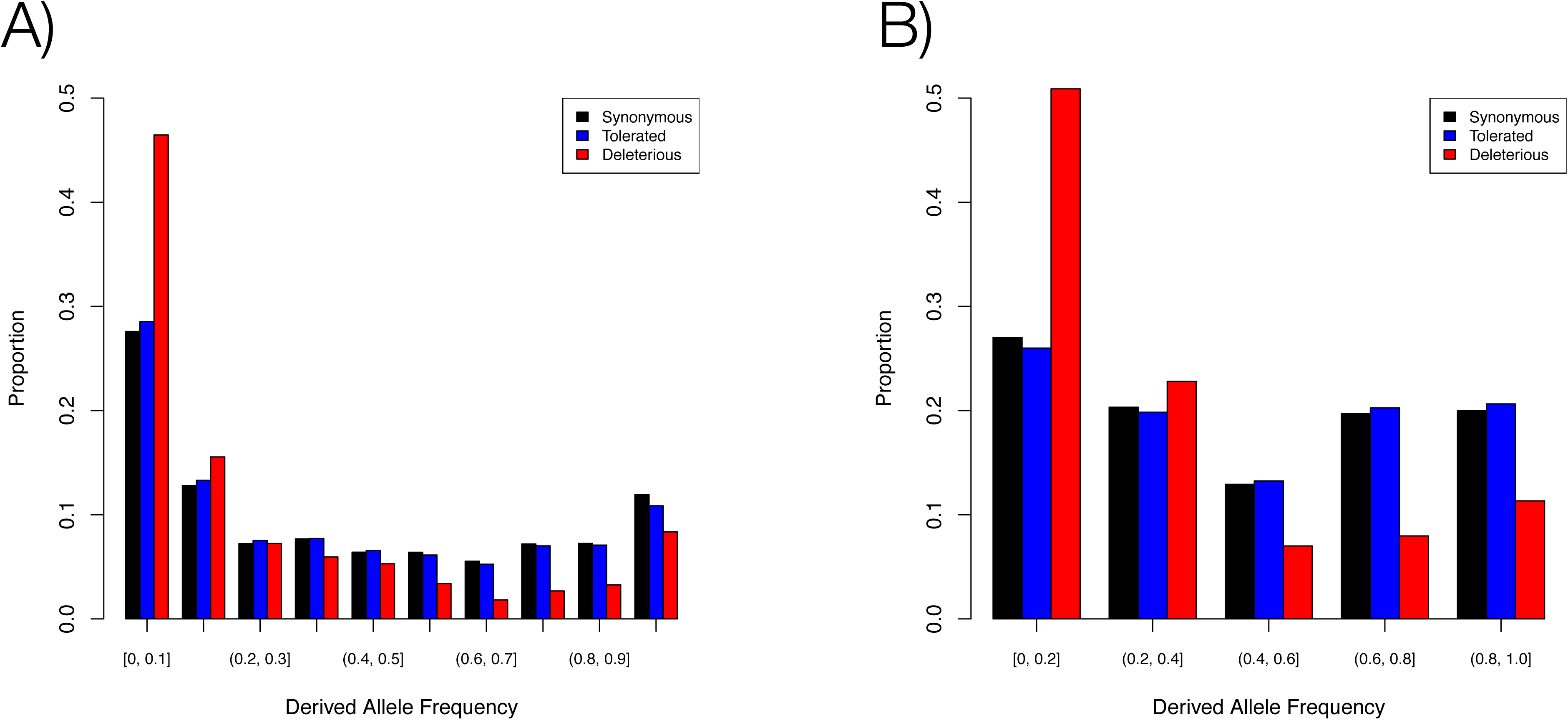
Derived allele (unfolded) frequency distributions for coding regions showing deleterious, tolerated, and synonymous SNPs for barley and soybean. Ancestral state was inferred as described in the methods. A variant was called “Deleterious” if it was nonsynonymous and predicted to be deleterious by SIFT, PolyPhen2, and the LRT. A) is based on thirteen domesticated barley accessions and two wild accessions while B) is based on seven cultivated soybean accessions and one wild accession.

In soybean, we called 586,102 SNPs in gene regions. Of these, 542,558 occured in the flanking regions of a gene model. We identified 73,577 synonymous SNPs, and 99,685 nonsynonymous SNPs (Table S5). SNPs in the various classes sum to greater than the total number of SNPs as a single SNP in multiple transcripts can have multiple functional annotations. For instance, a SNP may be intronic in one transcript, but exonic in an alternative transcript. SIFT identified 7,694 of the nonsynonymous SNPs as deleterious, PolyPhen2 identified 14,933 as deleterious, and the LRT identified 11,223 as deleterious. Per-sample counts of putatively deleterious variants in barley are shown in Table S6, and per-sample counts for soybean are shown in Table S7. Similarly to the barley sample, the proportion of putatively deleterious variants was similar across prediction approaches, with the exception of SIFT, which failed to find alignments for many genes. The overlap of prediction approaches identified 3,041 (2.6%) of nonsynonymous variants to be deleterious (Table 2). Derived allele frequency distributions are shown in Figure 1B. Variants inferred to be deleterious are generally at lower derived allele frequency than other classes of variation, implying that these variants are truly deleterious. For both species, the intersection of approaches appeared to give the most accurate prediction of whether or not a variant is deleterious, as evidenced by enrichment for rare alleles (Figure 2).

**Table 2.**
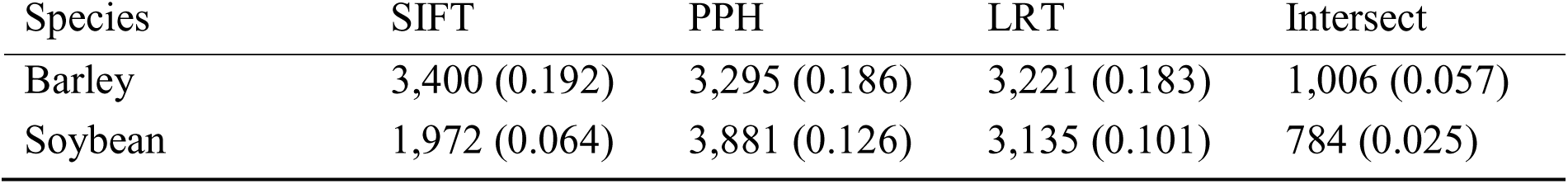
Mean counts of nonsynonymous variants that are predicted to be deleterious by three prediction methods. Numbers in parentheses are proportions of all nonsynonymous variants in each sample that are predicted to be deleterious.

**Figure 2:**
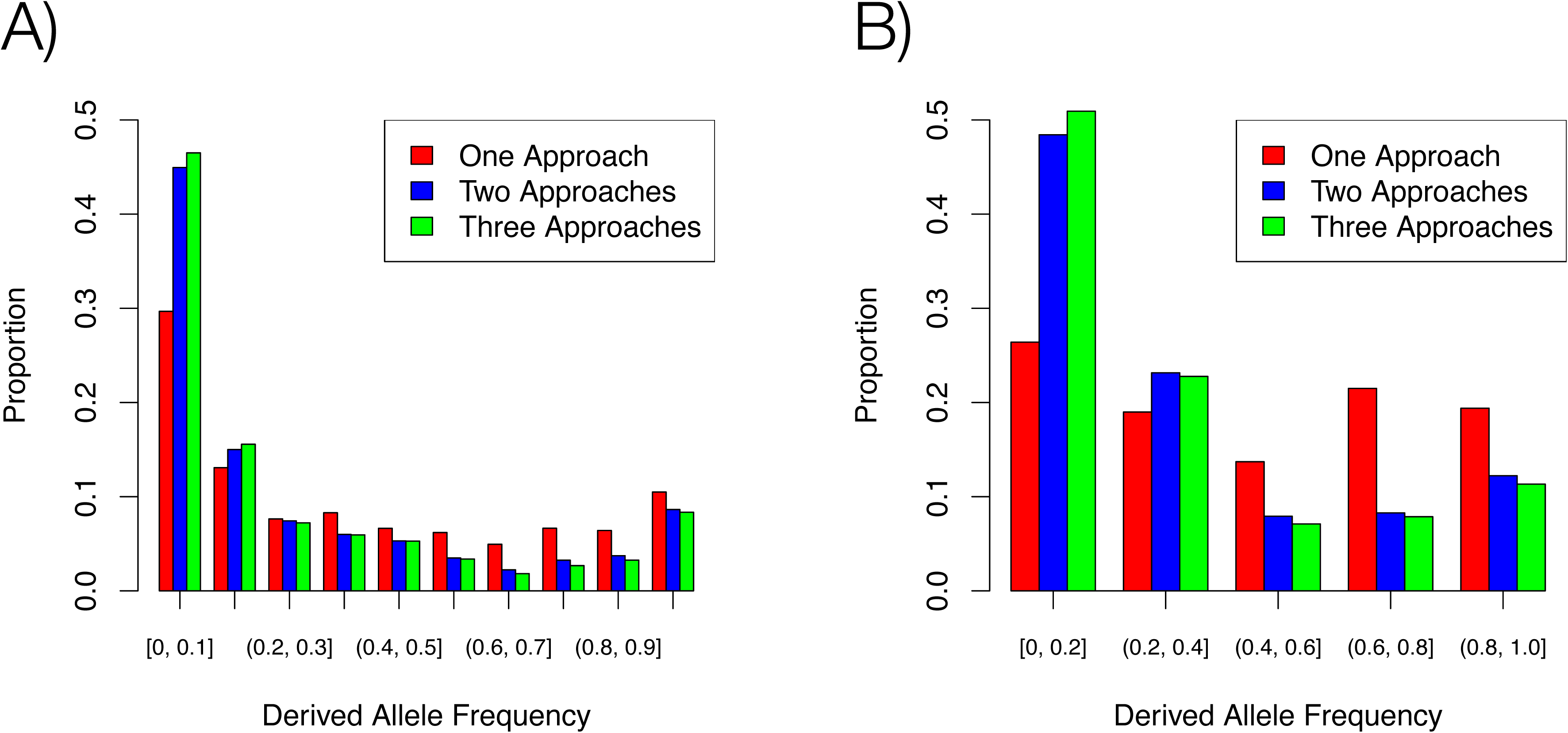
Derived allele (unfolded) frequency distributions for SNPs in A) barley and B) soybean predicted to be deleterious by one, two, or three prediction approaches. SNPs predicted by only one approach are not as strongly skewed toward rare variants, suggesting that the intersection of multiple prediction approaches gives the most reliable prediction of deleterious variants.

Nonsense variants made up a relatively small proportion of putatively deleterious variants. In our barley sample, we identified a total of 711 nonsense variants, 14.5% of our putatively deleterious variants. In soybean, we identified 1,081 nonsense variants, which make up 15.7% of putatively deleterious variants. Nonsense variants have a higher heterozygosity than tolerated, silent, or deleterious missense variants (Figure S1). While the absolute differences in heterozygosity were small due to the inbred nature of our samples, the pattern suggests that nonsense variants are more strongly deleterious than missense variants.

The transition to transversion ratio in our barley samples was 1.7:1 (Figure S2B), very close to estimates obtained from previous Sanger resequencing in barley genes (Morrell et al. 2006). In soybean, the transition to transversion ratio in our SNPs was 1.4:1, while the estimate from a Sanger resequencing dataset was ~1.2:1 (Hyten et al. 2006). The differences we observe between results from Sanger and Illumina resequencing could be due to the duplicated nature of the soybean genome (Schmutz et al. 2010), leading to paralogous alignment.

### Deleterious Mutations and Causative Variants

Bioinformatic approaches to identifying deleterious variants rely on sequence constraint to estimate protein functional impact. An example of a deleterious variant showing a derived base substitution that alters a phylogenetically conserved codon is shown in Figure S2. The variants identified in these approaches should be enriched for variants that cause large phenotypic changes. To explore how frequently known causative SNPs annotate as deleterious, we compiled a list of 23 nonsynonymous variants inferred to contribute to known phenotypic variation in barley and 11 in soybean, and tested the effect of these variants in our prediction pipeline. Of 23 variants that are purported to be causative for large phenotypic changes, 6 (25%) were inferred to be deleterious (Table S8). Of the 11 soybean putatively causative variants, 5 (45%) were inferred to be deleterious. This contrasts with the genome-wide average of ~3-6%, showing that variants our pipeline identifies as deleterious are more likely to impact phenotypes.

### Deleterious Mutations and Genetic Map Distance

The effective recombination rate strongly influences purging of deleterious variants from populations. To examine the relationship between the number of deleterious variants and recombination rate, we used a high-density genetic map from a soybean recombinant inbred line family (Lee et al. 2015). The soybean map was based on a subset of the SoySNP50K genotyping platform (Song et al. 2013). There was a weak but significant correlation between recombination rate and the proportion of nonsynonymous SNPs inferred to be deleterious (*r*^2^ = 0.007, p < 0.001, Figures 3, S3). We did not examine this relationship in barley because the barley reference genome assembly (Mayer et al. 2012) contains limited physical distance information.

**Figure 3:**
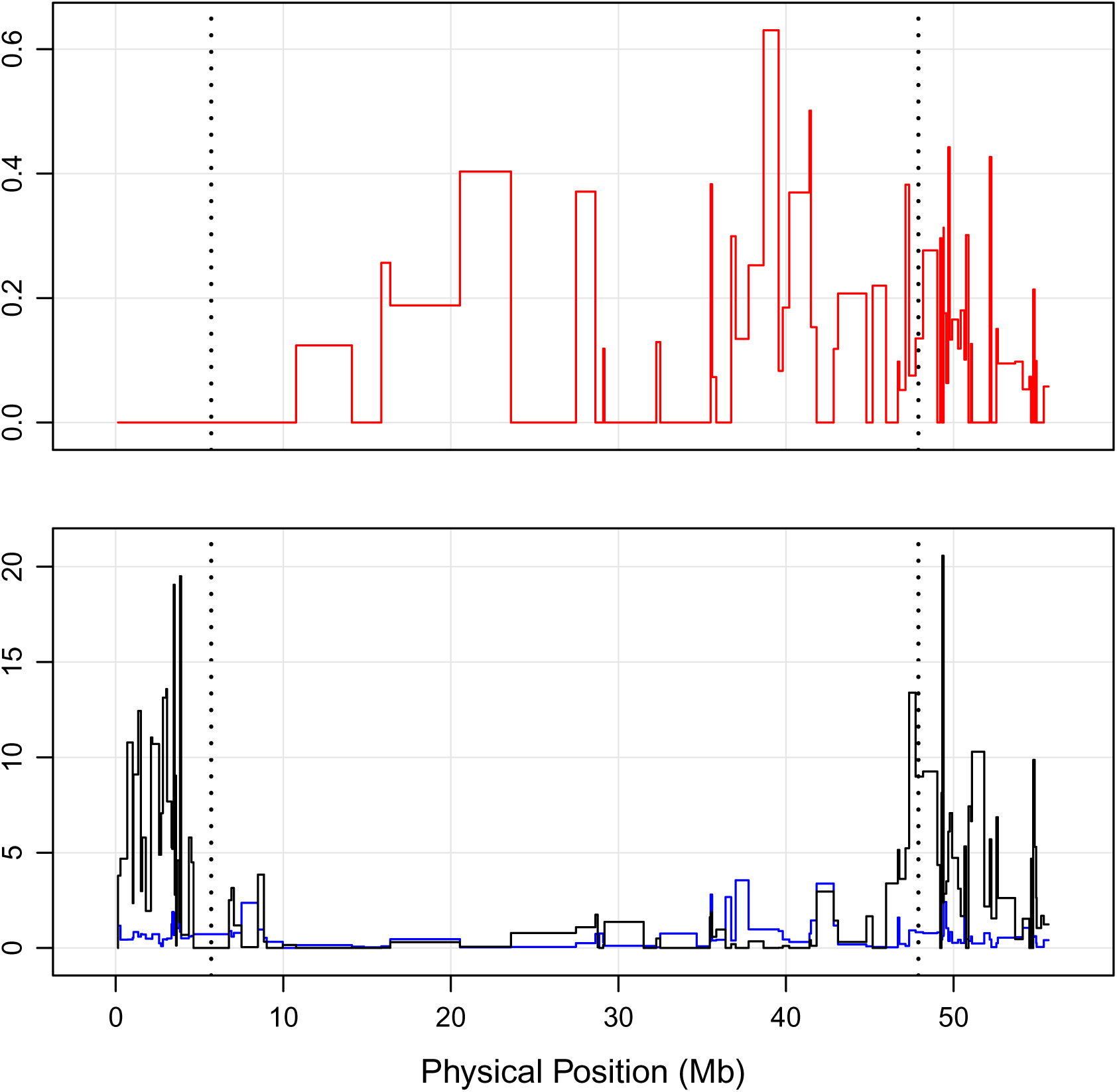
Comparison between recombination rate, coding sequence diversity, and proportion of nonsynonymous SNPs inferred to be deleterious in soybean on chromosome 1. The top panel shows the proportion of nonsynonymous SNPs that were inferred to be deleterious, in windows defined by genetic map distance (Lee et al. 2015). The bottom panel shows recombination rate in cM/Mb (black line) and average pairwise nucleotide sequence diversity per kilobase in coding sequence (blue line). Dashed vertical lines represent the boundaries of the annotated pericentromeric region, which has much lower recombination rates than the euochromatic regions.

## Discussion

Questions regarding the prevalence of deleterious variants date back over half a century (Fisher 1930; Muller 1950). In finite populations, the segregation of deleterious variants can have a substantial impact on population mean fitness (Kimura et al. 1963). While it has been argued that the concept of a reduction of fitness relative to a hypothetical optimal genotype is irrelevant (Wallace 1970), mutation accumulation studies have shown that new mutations have a substantial effect on absolute fitness (Schultz et al. 1999; Shaw et al. 2002).

Our results demonstrate that a large number of putatively deleterious variants persist in individual cultivars in both barley and soybean. The approaches used in this study predict the probability that a given amino acid or nucleotide substitution disrupts protein function. Mutations that alter phenotypes may be especially likely to annotate as deleterious, and we show that a high proportion of inferred causative mutations annotate as deleterious. It should be noted that variants identified as deleterious may affect a phenotype that is adaptive in only part of the species range or has a transient selective advantage – i.e., locally or temporally adaptive phenotypes. Our panel of causative variants consists primarily of SNPs that confer an agronomically important phenotype (Table S5). Agronomic phenotypes may be beneficial in wild populations, particularly biotic and abiotic stress tolerance or reproductive traits (Mercer et al. 2007), but are not expected to be either globally deleterious or globally beneficial. If the portion of the range in which the phenotype is adaptive is small or the selective advantage is transient, such variants will be kept at low frequencies and also be identified as deleterious. Just as few variants are expected to be globally advantageous, a portion of deleterious variation is likely to not be globally deleterious. Such variants could be either locally or temporally advantageous, with a fitness advantage under some circumstances contributing to their maintenance in populations (Tiffin and Ross-Ibarra 2014).

At the molecular level, variants occurring in minor transcripts of genes may exhibit conditional neutrality (Tiffin and Ross-Ibarra 2014), and *N_e_s* will be too low for purifying selection to act. Gan et al. (2011) identified many isoforms of genes among a diverse panel of *Arabidopsis thaliana* accessions, as well as compensatory mutations for a majority of frameshift mutations. Genetic variants that annotated as nonsynonymous or nonsense using the *A. thaliana* reference are frequently spliced out of the transcript such that the gene still produces a full-length and functional product. In a similar vein, deleterious variants may have their fitness impacts offset by compensatory mutations. In a study of bacteriophage, approximately 70% of deleterious mutations were offset by compensatory mutations (Poon and Chao 2005). The occurrence of the bulk of putatively deleterious variants in the lowest frequency classes (Figure 1), and a higher level of observed heterozygosity for putatively deleterious variants (Figure S1) are both consistent with the action of purifying selection on variants with negative impacts on fitness. Putatively disease-causing variants in human populations have also been observed to occur at low frequencies and to occur over a more geographically restricted range (Marth et al. 2011).

Identifying variants with low minor allele frequency (MAF) is an inexorable part of studying variants with fitness impacts. This presents a problem, as rare variants are the most likely to be affected by false positive variant calls, as they are necessarily observed very few times in the sample. In an attempt to abate the problem of false positive variant calls, we took an iterative approach to variant calling, applying strict genotype quality, read depth, and observed heterozygosity filters to reduce raw variant calls to a high-confidence set of variants. While it is true that some of the variants in our high-confidence set are false positives, they do not dominate our dataset. This is evidenced in our site frequency spectra (Figure 1), which does not indicate a strong skewing of putatively neutral variants toward low frequency classes, a pattern expected of genotyping errors. Additionally, false positive variants are expected to occur randomly, which would lead to roughly equal numbers of first, second, and third position SNPs within codons. Our variant calls show a strong enrichment toward third positions in codons (Figure S2), which are mostly synonymous positions, and are expected to be neutral. Deficiencies in first and second positions, which are mostly nonsynonymous sites under purifying selection, are indicative of our variant calls consisting mostly of true positive variants.

### Comparison of Identification Methods

Each of the approaches used here to identify deleterious variants makes use of sequence constraint across a phylogenetic relationship. They differ in terms of the models used to assess the functional effect of a variant. SIFT uses a heuristic, which determines if a nonsynonymous variant alters a conserved site based on an alignment build from PSI-BLAST results (Ng 2003). Polyphen2 is similar, but additionally identifies potential disruptions in secondary or tertiary structure of the encoded protein (when this information is available) (Adzhubei et al. 2010), and is trained on known human disease-causing polymorphisms and neutral polymorphisms. Both of these approaches estimate codon conservation from a multiple sequence alignment, but do not use phylogenetic relationships in their predictions. PolyPhen2 identified the largest number of variants as deleterious, perhaps reflecting bias from the training dataset. Nonhuman systems may differ fundamentally as to which amino acid substitutions tend to have strong functional impact, which would reduce prediction accuracy in other species (Adzhubei et al. 2010). The LRT explicitly calculates the local synonymous substitution rate, and uses it to test whether an individual codon is under selective constraint or evolving neutrally (Chun and Fay 2009). It is a hypothesis-driven approach, and compares the likelihood of two evolutionary scenarios. Variants in selectively constrained codons are considered to be deleterious.

Our results show that even though each prediction approach identifies a similar proportion of nonsynonymous SNPs as deleterious, the overlap between approaches is very small. Because each approach varies slightly in its prediction procedure and assumptions, the intersection of multiple approaches may provide more accurate predictions than any single prediction approach alone. At the genome-wide scale, this pattern is apparent in the frequency distribution of the variants that are identified as deleterious by all three approaches. They are enriched in the lowest frequency class, suggesting that they are under purifying selection (Figure 2). Further, known phenotype-altering SNPs are more likely to be predicted to be deleterious by all three approaches than those without known or measurable phenotypic impacts. This suggests that the intersection of prediction approaches tends to identify variants that are more likely to have fitness consequences, especially if the variant has a large effect on a phenotype. Identifying variants that are likely to have large effects on protein function and phenotype improves our ability to identify the nature of trait variation, especially if rare alleles of large effect are major contributors to complex traits (Thornton et al. 2013).

The SNPs predicted to be deleterious differ somewhat between prediction approaches. Even though SIFT and PolyPhen2 identify similar proportions of nonsynonymous SNPs as deleterious, they overlap at only approximately 50% of sites (Table 2). SNPs identified through at least two approaches seem more likely to be deleterious, based on lower average derived allele frequencies (Figure 1). Comparisons of the distribution of Grantham scores (Grantham 1974) show high similarity in the severity of amino acid replacements that are predicted to be deleterious by each approach (Figure S4). The effects of reference bias are apparent in SIFT and PolyPhen2. In barley and soybean, the reference genotypes are 'Morex' and ‘Williams 82’ respectively. Even when polarized by ancestral and derived alleles, these genotypes show considerably fewer inferred deleterious variants (Table S6; Table S7).

The software package we have developed to implement the LRT is called BAD_Mutations (<u>B</u>LAST <u>A</u>ligned-<u>D</u>eleterious Mutations;). While BAD_Mutations is similar in approach to SIFT and PolyPhen2, it uses distinct data sources and models to predict whether or not a SNP is deleterious. SIFT and PolyPhen2 rely on BLAST searches against a general nucleotide sequence database, which results in high degree of variability in data quality from gene to gene (data not shown). BAD_Mutations, on the other hand, uses a set of assembled and annotated genome sequences available in the public domain in databases such as Phytozome (https://phytozome.jgi.doe.gov) and Ensembl Plants (http://plants.ensembl.org/). The use of a standard set of genome sequences helps to ensure consistent phylogenetic comparisons for each gene analyzed. It also uses a model that weights the conservation of the amino acid residue by the synonymous substitution rate of the gene under consideration (Chun and Fay 2009). BAD_Mutations is open source and freely available at 

~~~
https://github.com/MorrellLAB/BAD_Mutations
~~~

.

### Rise of Deleterious Variants Into Populations

The number of segregating deleterious variants in a species is very different from the number of *de novo* deleterious mutations in each generation, commonly identified as *U. U*is the product of the per-base pair mutation rate, the genome size, and the fraction of the genome that is deleterious when mutated (Charlesworth 2012). In humans, *U* is estimated at ~2 new deleterious variants per genome per generation (Agrawal and Whitlock 2012). Estimates from *Arabidopsis thaliana* suggest that the genomic mutation rate for fitness-related traits is ~0.1-0.2 per generation (Shaw et al. 2002), and approximately half are estimated to be detrimental to fitness. Even though new mutations are constantly arising, the standing load of deleterious variation greatly exceeds the rate at which they arise (Charlesworth et al. 2004; Charlesworth 2012). However, our results show that ~40% of our inferred deleterious variants are private to individual cultivars, suggesting that they can be purged from breeding programs.

Once deleterious variants are present as segregating variation in the progenitors of crops, genetic bottlenecks associated with domestication (Eyre-Walker et al. 1998) may allow deleterious variants to drift to higher frequency (Robertson 1960). The selective sweeps associated with domestication and improvement (Wright et al. 2005) would decrease nucleotide diversity in affected genomic regions (Smith and Haigh 1974; Kaplan et al. 1989), and subsequently reduce the effective recombination rate (cf. O’Reilly 2008). The selective and demographic processes of domestication and improvement lead to three basic hypotheses about the distribution of deleterious variants in crop plants: *i*) the relative proportion of deleterious variants will be higher in domesticates than in wild relatives; *ii*) deleterious variants will be enriched near loci of agronomic importance that are subjected to strong selection during domestication and improvement; *iii*) the relative proportion of deleterious variants will be lower in elite cultivars than landraces due to strong selection for yield (Gaut et al. 2015). Future studies of deleterious variants in crops and their wild relatives can address these hypotheses to understand the source of variation in modern cultivated material.

### Deleterious Variants in Crop Breeding

The identification and targeted elimination of deleterious variants has been proposed as a potential means of improving plant fitness and crop yield (Morrell et al. 2011). Current plant breeding strategies using genomewide prediction rely on estimating genome-wide marker effects on quantitative traits of interest (Meuwissen et al. 2001). Genomewide prediction has been shown to be effective in both animals (Schaeffer 2006) and plants (Heffner et al. 2011; Jacobson et al. 2014), but these approaches rely on estimating marker contributions to a quantitative trait (i.e., a measured phenotypic effect). The genetic architecture of these traits suggests that our ability to quantify the effects of individual loci will reach practical limits before we can identify loci contributing to their variance (Rockman 2012). QTL mapping approaches to identifying favorable variants for agronomic traits will reach practical limits, even for variants of large effect (King et al. 2012). Many traits of agronomic interest, particularly yield in grain crops, are quantitative and have a complex genetic basis. As such, they are under the influence of environmental effects and many loci (Falconer and Mackay 1996). Current genomewide prediction and selection methodologies rely on estimating the combined effects of markers across the genome (Meuwissen et al. 2001), but this approach is limited by recombination rate and the ability to measure phenotypes of interest. The identification and purging of deleterious variants should provide a complementary approach to current breeding methodologies, if bioinformatically-identified deleterious variants are truly deleterious (Morrell et al. 2011).

In the current study, we restricted our analyses to protein coding regions, but additional recent evidence suggests that deleterious variants can accumulate in conserved noncoding sequences, such as transcription factor binding sites (Arbiza et al. 2013). Additionally, insertion and deletion polymorphisms and larger structural variants were not considered in this study. Structural variants are abundant in crop plants, and may be involved with large phenotypic changes (Chia et al. 2012; Anderson et al. 2014). As such, analysis of nonsynonymous SNPs presents a lower bound on the estimates of the number of deleterious variants segregating in populations. Efforts to identify deleterious variants in noncoding sequence are limited by scant knowledge of functional constraints on noncoding genomic regions, and difficulty in aligning noncoding regions from all but the most closely related taxa (Doniger et al. 2008). Annotation of noncoding sequence will uncover additional deleterious variants, but the roughly one thousand putatively deleterious variants we identify per individual cultivar should provide ample targets for selection of recombinant progeny in a breeding program.

## Materials and Methods

### Plant Material and DNA Sequencing

The exome resequencing data reported here includes thirteen cultivated barleys, and two wild barley accessions. Barley exome capture was based on a 60 Mb liquid-phase Nimblegen capture design (Mascher et al. 2013). For the soybean sample, we resequenced whole genomes of seven soybean cultivars and used previously-generated whole genome sequence of *Glycine soja* (Kim et al. 2010). Each sample was prepared and sequenced with manufacturer protocols (Illumina, San Diego, CA) to at least 25x coverage of the target with 76bp, 100bp or 151bp paired-end reads. A summary of samples and sequencing statistics is given in Table S1.

### Read Mapping and SNP Calling

DNA sequence handling followed the “Genome Analysis Tool Kit (GATK) Best Practices” workflow from the Broad Institute (McKenna et al. 2010; DePristo et al. 2011). Our workflow for read mapping and SNP calling is depicted in Figure S1. First, reads were checked for proper length, Phred score distribution, and *k*-mer contamination with FastQC (bioinformatics.babraham.ac.uk/projects/fastqc/). Primer and adapter sequence contamination was then trimmed from barley reads using Scythe (github.com/vsbuffalo/scythe), using a prior on contamination rate of 0.05. Low-quality bases were then removed with Sickle (github.com/najoshi/sickle), with a minimum average window Phred quality of 25, and window size of 10% of the read length. Soybean reads were trimmed using the fastqc-mcf tool in the eautils package (code.google.com/p/ea-utils/). Post-alignment processing and SNP calling were performed with the GATK v. 3.1 (McKenna et al. 2010; DePristo et al. 2011).

Barley reads were aligned to the Morex draft genome sequence (Mayer et al. 2012) using BWA-MEM (Li and Durbin 2009). We tuned the alignment reporting parameter and the gapping parameters to allow ~2% mismatch between the reads and reference sequence, which is roughly equivalent to the highest estimated nucleotide diversity observed at a locus in barley coding sequence (Morrell et al. 2003, 2006, 2014). The resulting SAM file was trimmed of unmapped reads with Samtools (Li et al. 2009), sorted, and trimmed of duplicate reads with Picard tools (picard.sourceforge.net/). We then realigned around indels, using a set of 100 previously known indels from Sanger resequencing of 25 loci (Caldwell et al. 2006; Morrell and Clegg 2007; Morrell et al. 2014). Sequence coverage was estimated with ‘bedtools genomecov,’ using the regions included in the Nimblegen barley exome capture design (https://sftp.rch.cm/diagnostics/sequencing/nimblegen_annotations/ez_barley_exome/barley_exome.zip). Individual sample alignments were then merged into a multisample alignment for variant calling. A preliminary set of variants was called with the GATK HaplotypeCaller with a heterozygosity (average pairwise diversity) value of 0.008, based on average coding sequence diversity reported for cultivated barley (Morrell et al. 2014). This preliminary set of variants was filtered to sites with a genotype score of 40 or greater, heterozygous calls in at most two individuals, and read depth of at least five reads. We then used the filtered variants, SNPs identified in the Sanger resequencing data set, and 9,605 SNPs from genotyping assays: 5,010 from the James Hutton Institute (Comadran et al. 2012), and 4,595 from Illumina GoldenGate assays (Close et al. 2009) as input for the GATK VariantRecalibrator to obtain a set of recalibrated variant calls. Final variants were filtered to be supported by a minimum of five reads per sample, have a Phredscaled genotype quality of at least 40, and have a maximum of two accessions with missing data.

Processing of soybean samples is as described above, but with the following modifications. Soybean reads were aligned to the Williams 82 reference genome sequence (Schmutz et al. 2010). Mismatch and reporting parameters for the cultivated samples were adjusted to allow for ~1% mismatch between reads and reference, which is approximately the highest coding sequence diversity typically observed in soybean (Hyten et al. 2006). The alignments were trimmed and sorted as described above. Preliminary variants were called as in the barley sample, but with a heterozygosity value of 0.001, which is the average nucleotide diversity reported by Hyten et al. (2006). Final variant calls were obtained in the same way as described for the barley sample, using SNPs on the SoySNP50K chip (Song et al. 2013) as known variants.

Transition to transversion ratios were calculated with R scripts. The ratios in the Sanger resequencing dataset were computed using SNPs identified in FASTA alignments of wild barley gene sequences (Morrell et al. 2006), or a table of SNPs identified in resequencing of soybean gene fragments (supplemental data file 1 in Hyten et al. 2006).

Read mapping scripts, variant calling scripts, and variant filtering scripts for both barley and soybean are available on GitHub at (github.com/MorrellLAB/Deleterious_Mutations).

### SNP Classification

Barley SNPs were identified as coding or noncoding using the Generic Feature Format v3 (GFF) file provided with the reference genome (Mayer et al. 2012). A custom Python script was then used to identify coding barley SNPs as synonymous or nonsynonymous. Soybean SNPs were assigned using primary transcripts using the Variant Effect Predictor (VEP) from Ensembl (ensembl.org/info/docs/tools/vep/index.html). Nonsynonymous SNPs were then assessed using SIFT (Ng 2003), PolyPhen2 (Adzhubei et al. 2010) using the ‘HumDiv’ model, and a likelihood ratio test comparing codon evolution under selective constraint to neutral evolution (Chun and Fay 2009). For the likelihood ratio test, we used the phylogenetic relationships between 37 Angiosperm species based on genic sequence from complete plant genome sequences available through Phytozome (phytozome.jgi.doe.gov/) and Ensembl Plants (plants.ensembl.org/). The LRT is implemented as a Python package we call ‘BAD_Mutations’ (<u>B</u>LAST <u>A</u>ligned-<u>D</u>eleterious Mutations; github.com/MorrellLAB/BAD_Mutations). Coding sequences from each genome were downloaded and converted into BLAST databases. The coding sequence from the query species was used to identify the best match from each species using TBLASTX. The best match from each species was then aligned using PASTA (Mirab et al. 2014), a phylogeny-aware alignment tool. The resulting alignment was then used as input to the likelihood ratio test for the affected codon. The LRT was performed on codons with a minimum of 10 species represented in the alignment at the queried codon. Reference sequences were masked from the alignment to reduce the effect of reference bias (Simons et al. 2014). A SNP was identified as deleterious if the p-value for the test was less than 0.05, with a Bonferroni correction applied based on the number of tested codons, and if either the alternate or reference allele was not seen in any of the other species. For barley, our threshold was 8.4E-7 (59,277 codons tested), and for soybean, our threshold was 7.8E-7 (64,087 codons tested). A full list of species names and genome assembly and annotation versions used is available in Table S9.

### Relating Recombination Rate to Deleterious Predictions

Recombination rates were taken from a genetic map developed by Lee et al. (2015). Briefly, a recombinant inbred line family was derived from a cross between a wild soybean line and a cultivated soybean line, and genotyped with the SoySNP6K genotyping platform. For our analysis, we calculated cM/Mb values for each interval between markers on the SoySNP6K. Within each interval, we also calculated the proportion of nonsynonymous SNPs that annotated as deleterious by our criteria. Intervals with negative, or cM/Mb values above 20 were excluded, as they indicate regions where the markers likely have incorrect physical position. Pearson correlation (Figure S3A) and logistic regress (Figure S3B) were used to investigate the relationship between recombination rate and deleterious variation.

### Inference of Ancestral State

Prediction of deleterious mutations is complicated by reference bias (Chun and Fay 2009; Simons et al. 2014), which manifests in two ways. First, individuals that are closely related to the reference line used for the reference genome will appear to have fewer genetic variants, and thus fewer inferred nonsynonymous and deleterious variants. Second, when the reference strain carries a derived allele at a polymorphic site, that site is generally not predicted to be deleterious (Simons et al. 2014). To address the issue of reference bias, we polarized all coding variants by ancestral and derived state, rather than reference and non-reference state. Ancestral states were inferred for SNPs in gene regions by inferring the majority state in the most closely related clade from the consensus phylogenetic tree for the species included in the LRT. For barley, the ancestral states were inferred from gene alignments of *Aegilops tauschii, Brachypodium distachyon*, and *Tritium urartu*. For soybean, ancestral states were inferred using *Medicago truncatula* and *Phaseolus vulgaris*. This approach precludes universal inference of ancestral state for noncoding variants. However, examination of alignments of intergenic sequence in Triticeae species and in *Glycine* species showed that alignments outside of protein coding sequence is not reliable for ancestral state inference (data not shown).

## Acknowledgements

The authors thank Brandon S. Gaut, Michael B. Kantar, Ana M. Poets, and Ruth G. Shaw for helpful comments on an earlier version of the manuscript. The authors also thank two anonymous reviewers for their critique of the analysis and presentation. This work was supported by a USDA NIFA National Needs Fellowship (Appropriation No. 5430-21000-006-00D) and a MnDrive 2014 Food Security Fellowship, in support of TJYK. Support was also provided by the Minnesota Agricultural Experiment Station Variety Development fund, the United Soybean Board and U.S. NSF Plant Genome Program (DBI-1339393). This research was carried out with hardware and software support provided by the Minnesota Supercomputing Institute (MSI) at the University of Minnesota.

## Author Contributions

TJYK, RMS, PT, and PLM designed the research. KPS and PLM provided input on which barley lines to sample, and RMS provided sequence data for soybean lines. Barley read mapping, variant calling and assessment with SIFT and PolyPhen2 were performed by TJYK and CL. Soybean data analysis was performed by FF with assistance from TJYK. Code for the likelihood ratio test was developed by TJYK, PJH, and JCF. Breeding history and causative mutations list were provided by MM. TJYK and PLM wrote the manuscript.

## Table Captions

Table 1: Mean numbers of SNPs in various annotation classes in barley and soybean. The numbers in this table have not been polarized by ancestral or derived state.

Table 2: Mean numbers of nonsynonymous SNPs that are identified as deleterious by each of the prediction approaches, in barley and soybean. Numbers in parentheses are proportions of all nonsynonymous variants that annotate deleterious. Each approach identifies a similar number of deleterious SNPs, but the sets are mostly non-overlapping.

Table S1: Accessions used in this study. The final coverage reported is the average depth over the targeted region.

Table S2: Location and date of cultivar release for the barley accessions used in this study.

Table S3: Location and date of cultivar release for the soybean accessions used in this study.

Table S4: Counts of SNPs in various classes in thirteen barley accessions and two wild barley accessions. Numbers reported are comparisons against the reference genome, which makes it possible to include noncoding variants, where ancestral state cannot be estimated unambiguously

Table S5: Counts of SNPs in various classes in seven soybean accessions and one wild soybean accessions. Numbers reported are comparisons against the reference genome, which makes it possible to include noncoding variants, where ancestral state cannot be estimated unambiguously.

Table S6: Per-approach and per-sample counts of deleterious variants for barley. Numbers reported are comparisons against ancestral state. The proportion of nonsynonymous variants that is inferred to be deleterious by each prediction approach in each accession is shown in parentheses

Table S7: Per-approach and per-sample of counts of deleterious variants in soybean. Numbers reported are comparisons against ancestral state. The proportion of nonsynonymous variants that is inferred to be deleterious by each prediction approach in each accession is shown in parentheses.

Table S8: List of cloned genes with SNPs causing phenotypic differences, and predictions for each SNP. Causative SNPs annotate as deleterious with a higher frequency than the genomic average.

Table S9: List of all species and genome assembly versions, annotation versions, and data sources for sequences used in the likelihood ratio test.

**Figure S1:**
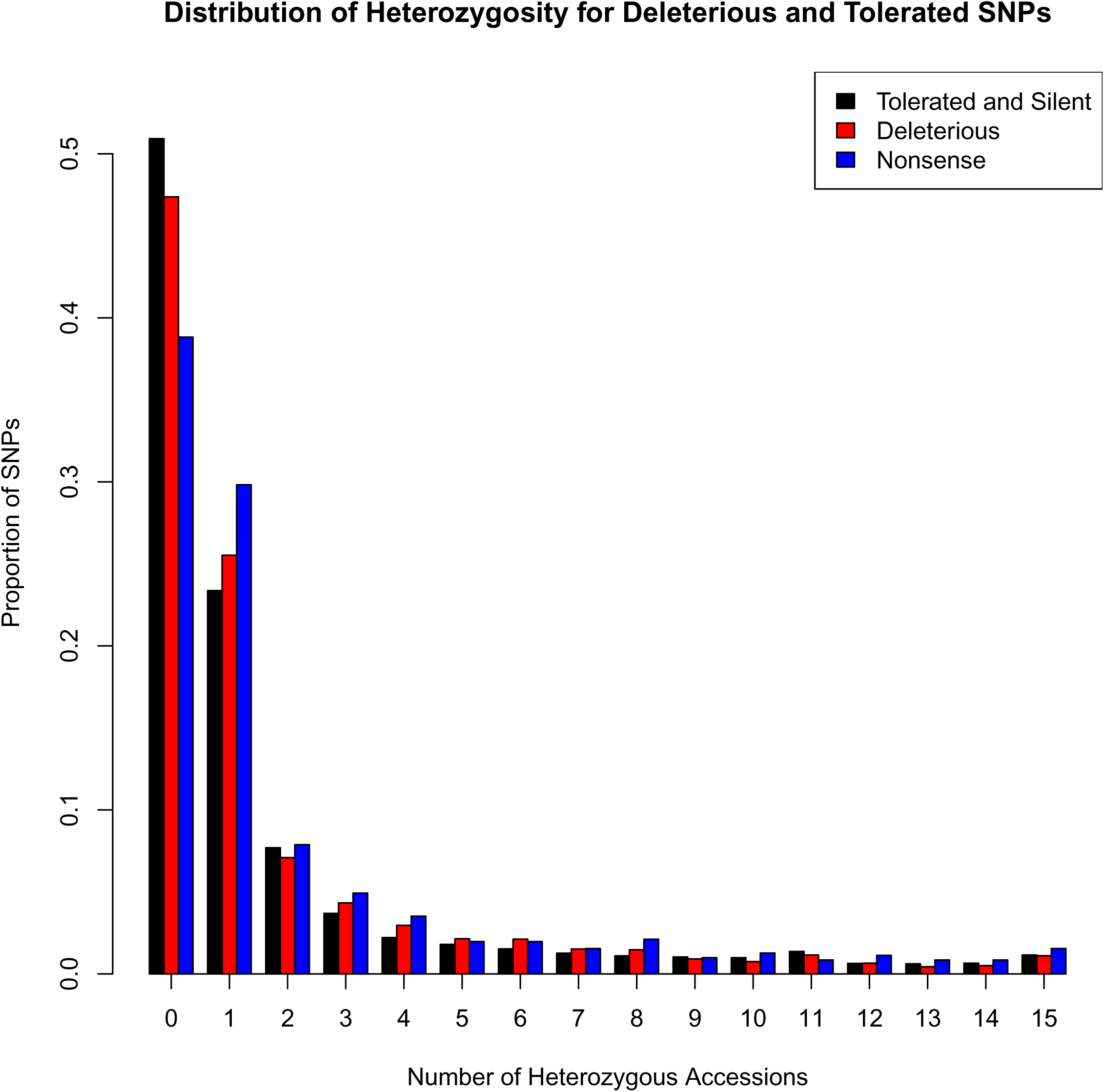
The distributions of per-SNP heterozygosity for tolerated nonsynonymous and silent SNPs, deleterious missense SNPs, and nonsense SNPs. Nonsense SNPs tend to be heterozygous more often than deleterious or tolerated SNPs.

**Figure S2:**
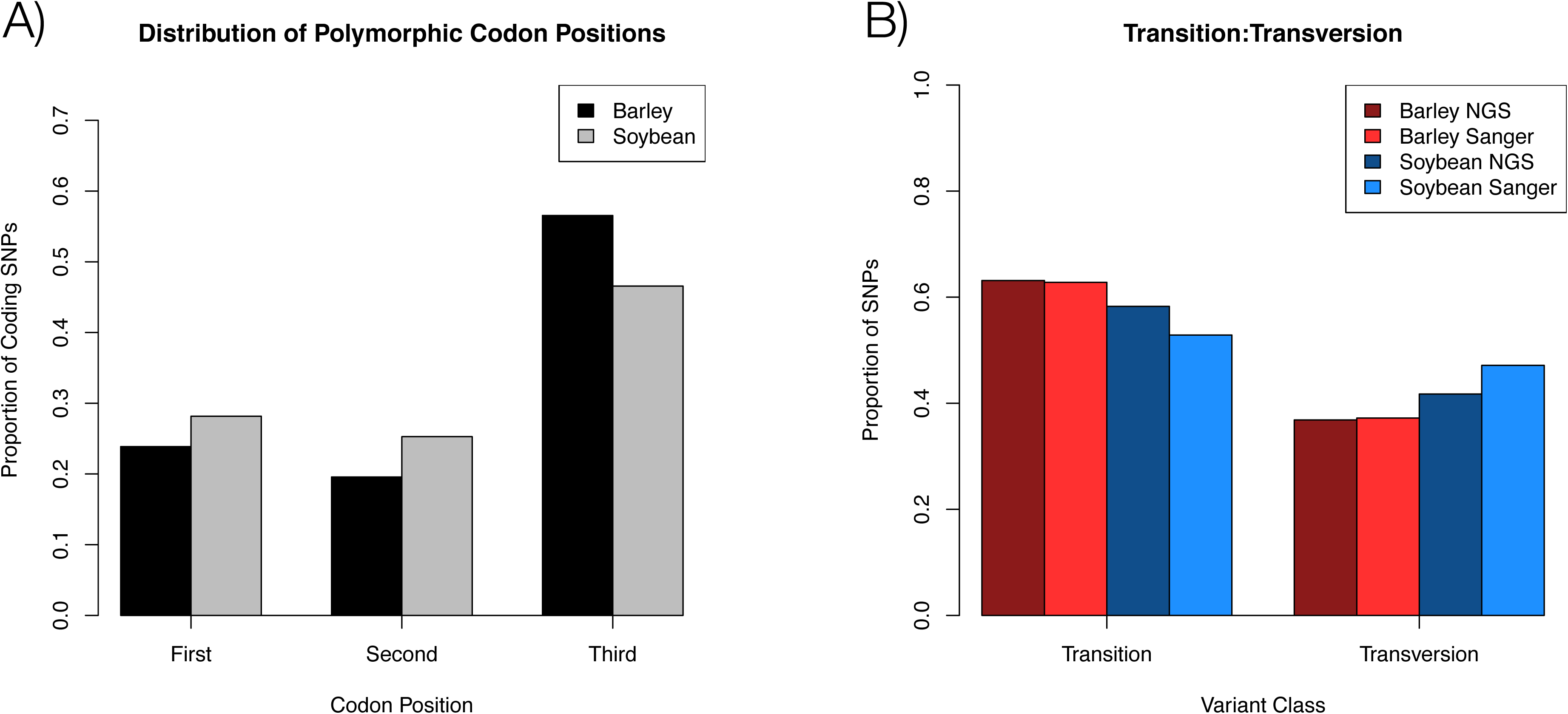
A) Histogram showing the relative frequencies of 1^st^, 2^nd^, and 3^rd^ position variants in codons. The distribution shows a strong bias against 1^st^ and 2^nd^ positions (which tend to be nonsynonymous), consistent with the action of purifying selection. B) Proportions of SNPs identified in our datasets that are transitions and transversions. The proportions estimated from Sanger resequencing come from Morrell et al. (2006) for barley, and Hyten et al. (2006) for soybean.

**Figure S3:**
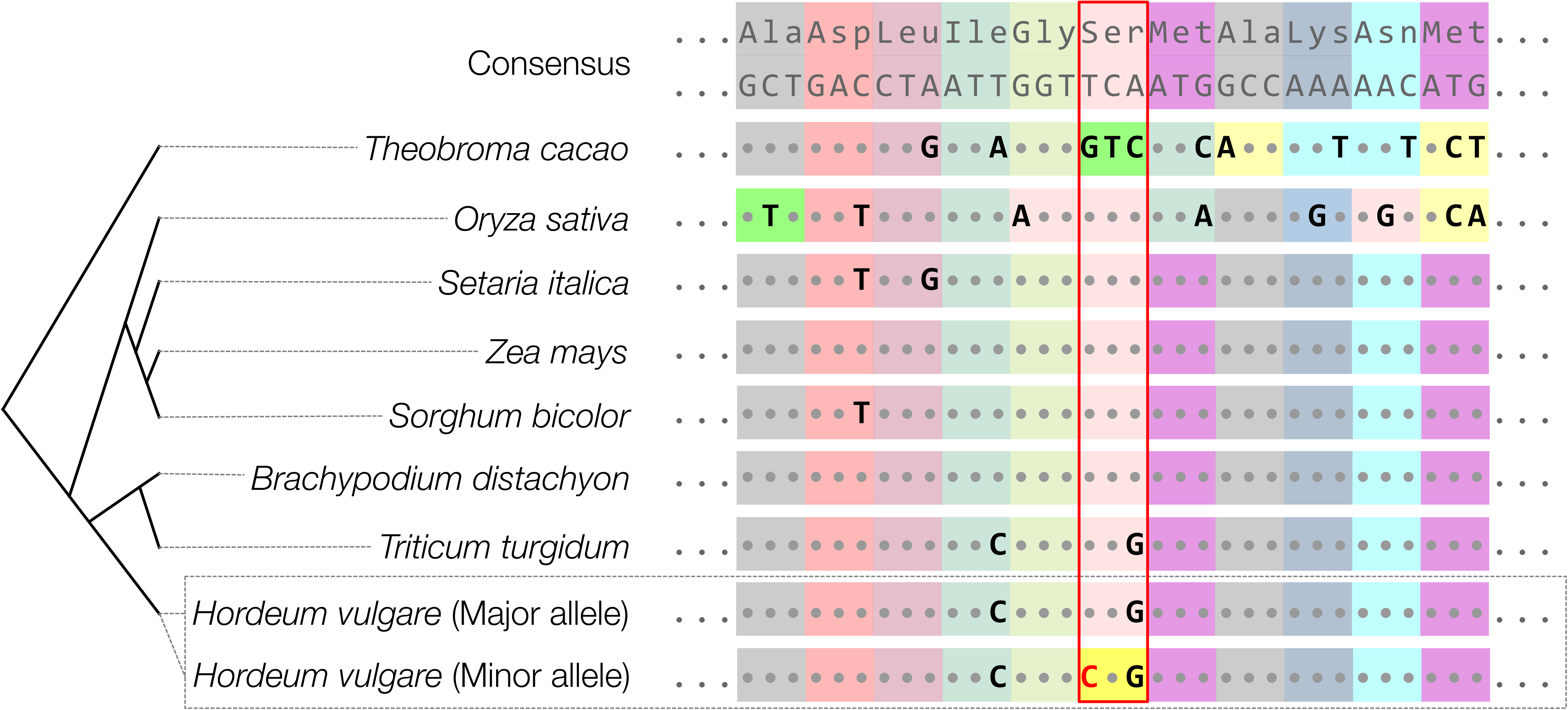
A sample alignment used to infer a Serine to Proline mutation as deleterious in *PpdH1*. The alignment is built from sequences used by SIFT, and the affected codon is highlighted in red.

**Figure S4:**
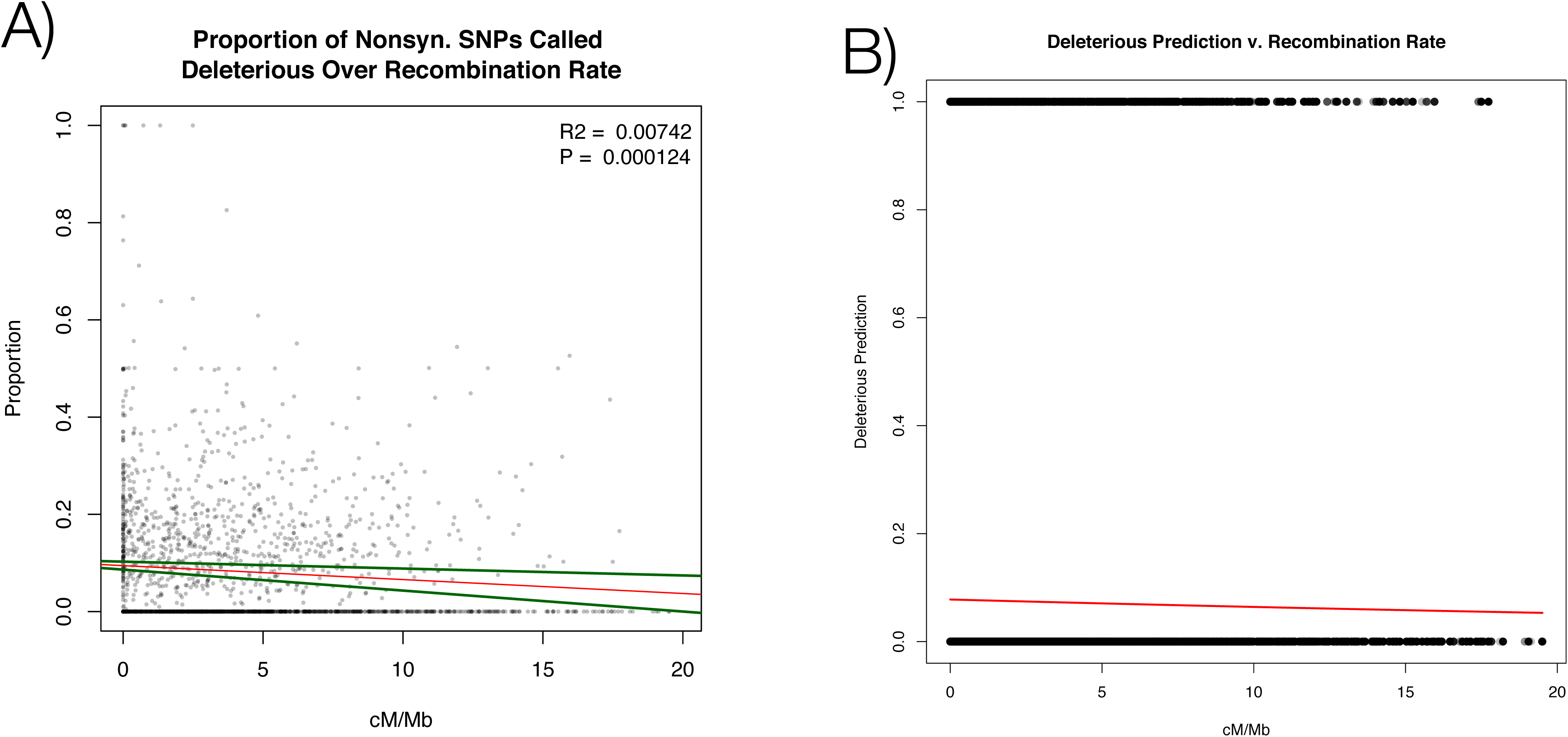
A) Correlation between recombination rate (cM/Mb) and proportion of nonsynonymous SNPs inferred to be deleterious genome-wide in our soybean sample. B) Logistic regression of whether or not a SNP is predicted to be deleterious against recombination rate in cM/Mb.

**Figure S5:**
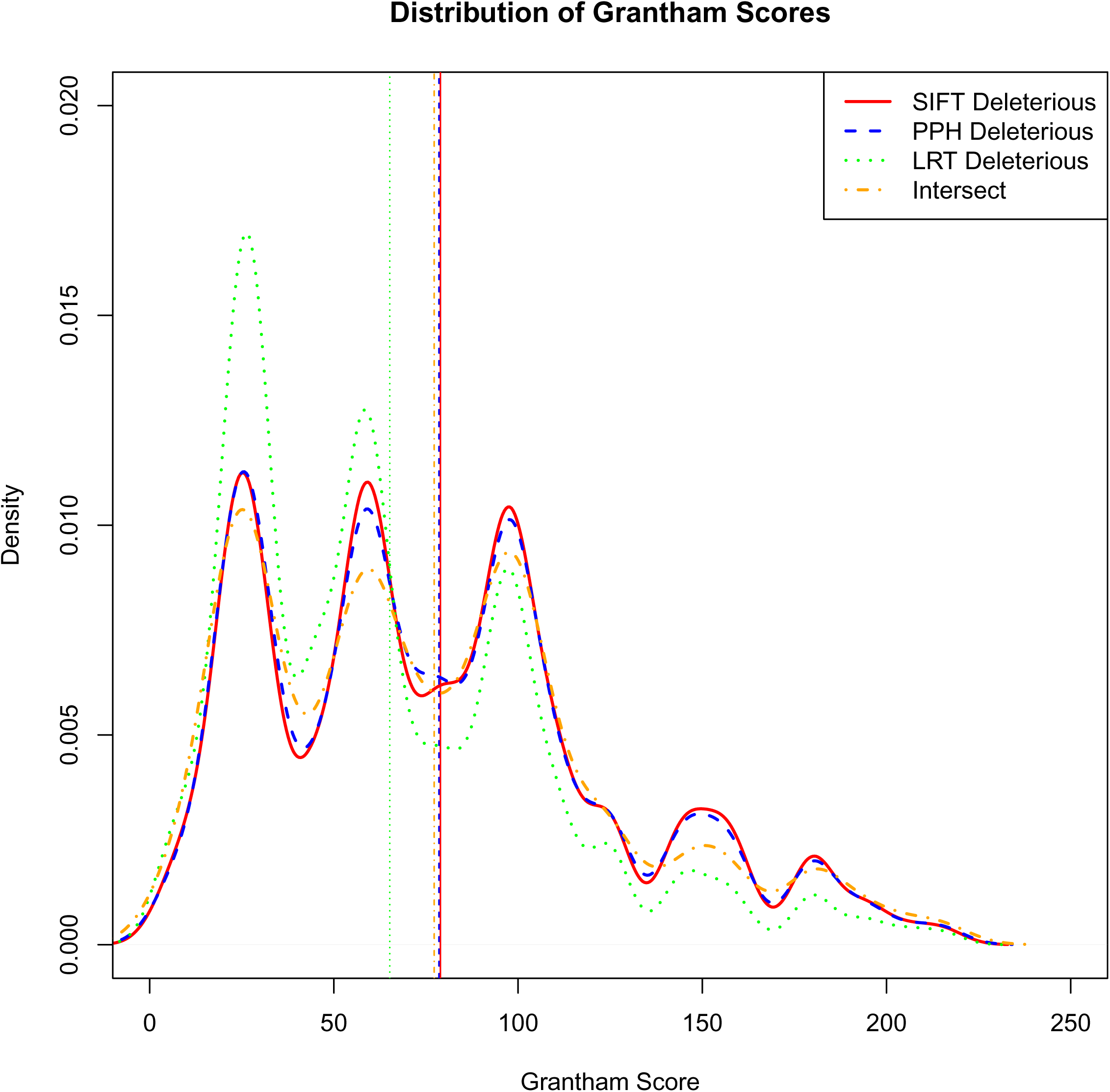
Distribution of Grantham score for nonsynonymous variants predicted to be deleterious by each prediction approach. Each approach and the intersection of each approach gives a very similar distribution of Grantham scores. Vertical lines show the mean of the distribution.

**Figure S6:**
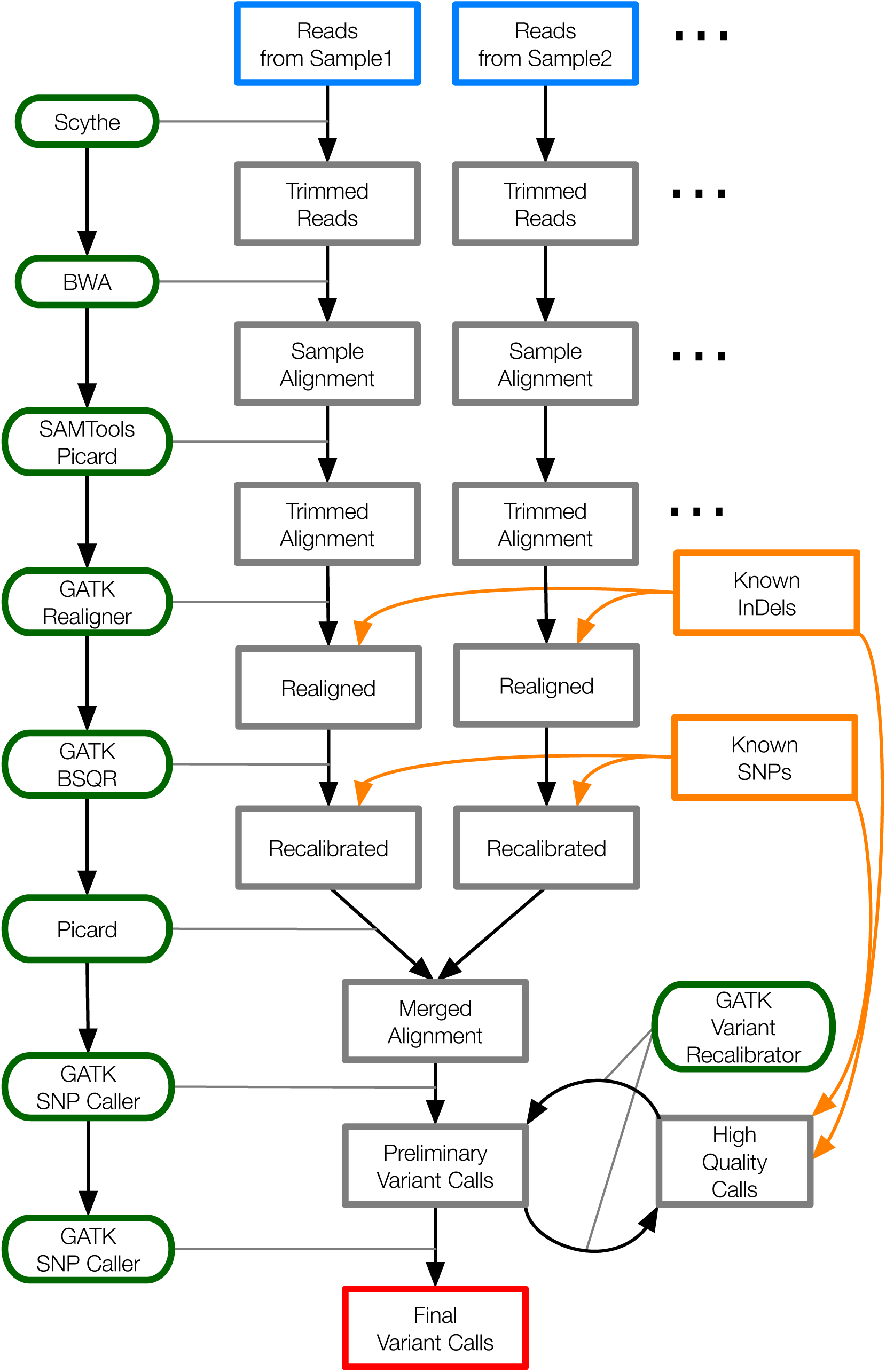
A schematic for the read mapping and SNP calling workflow. Boxes with bold borders denote the start and end points of workflow. Rounded boxes with light grey borders are the tools that are used at each step in the pipeline.

